# LACK OF EVIDENCE FOR A VIABLE MICROBIOTA IN MURINE AMNIOTIC FLUID

**DOI:** 10.1101/2021.08.10.455893

**Authors:** Andrew D. Winters, Roberto Romero, Jonathan M. Greenberg, Jose Galaz, Zachary Shaffer, Valeria Garcia-Flores, David J. Kracht, Nardhy Gomez-Lopez, Kevin R. Theis

## Abstract

The existence of an amniotic fluid microbiota (i.e., a viable microbial community) in mammals is controversial. Its existence would require a fundamental reconsideration of the role of intra-amniotic microbes in fetal development and pregnancy outcomes. In this study, we determined whether the amniotic fluid of mice harbors a microbiota in late gestation. Bacterial profiles of amniotic fluids located proximally or distally to the cervix were characterized through quantitative real-time PCR, 16S rRNA gene sequencing, and culture (N = 21 mice). These profiles were compared to those of technical controls for background DNA contamination. The load of 16S rDNA in the amniotic fluid exceeded that in controls. Additionally, the 16S rDNA profiles of the amniotic fluid differed from those of controls, with *Corynebacterium tuberculostearicum* being differentially more abundant in amniotic fluid profiles; however, this bacterium was not cultured. Of the 42 total bacterial cultures of amniotic fluids, only one yielded bacterial growth – *Lactobacillus murinus*. The 16S rRNA gene of this common murine-associated bacterium was not detected in any amniotic fluid sample, suggesting it did not originate from the amniotic fluid. No differences in 16S rDNA load, 16S rDNA profile, or bacterial culture were observed between amniotic fluids located proximal and distal to the cervix. Collectively, these data show that, although there is a modest DNA signal of bacteria in murine amniotic fluid, there is no evidence that this signal represents a viable microbiota. These findings refute the proposed role of amniotic fluid as a source of microorganisms for *in utero* colonization.

**IMPORTANCE:** The prevailing paradigm in obstetrics has been the sterile womb hypothesis, which posits that fetuses are first colonized by microorganisms during labor and/or the vaginal delivery process. However, it has been suggested that fetuses are consistently colonized *in utero*. One proposed source of colonizers is the amniotic fluid surrounding the fetus. This concept has been derived primarily from investigations that relied on DNA sequencing. Due to the low microbial biomass of amniotic fluid, such studies are susceptible to influences of background DNA contamination. Additionally, even if there is a microbial DNA signature in amniotic fluid, this is not necessarily reflective of a resident microbiota that could colonize the mammalian fetus. In the current study, using multiple microbiologic approaches and incorporating technical controls for DNA contamination, we show that, although there is a low abundance bacterial DNA signal in amniotic fluid, this does not translate to the presence of viable bacteria.

## INTRODUCTION

The mammalian amniotic cavity is filled with a protective liquid (i.e., amniotic fluid) that surrounds the fetus throughout gestation. Indeed, the amniotic fluid is essential for fetal development and maturation [1, 2]. As such, the amniotic fluid is enriched with nutrients and growth factors [1, 3–5] and contains soluble (e.g. cytokines [6–27], anti-microbial molecules, etc. [28–33]) and cellular (e.g. innate and adaptive immune cells [34–40]) components that serve as an immunological barrier against invading pathogens. In clinical medicine, the amniotic fluid is utilized as a diagnostic tool for assessing intra-amniotic inflammation and/or infection [41–59], a condition that is strongly associated with obstetrical disease, the most detrimental of which is preterm birth [60]. Therefore, the presence of microorganisms in the amniotic fluid is associated with adverse maternal and neonatal outcomes [61–67], and the traditional view in obstetrics has been the “sterile womb hypothesis”, which proposes that the fetal environment is sterile and that the neonate first acquires a microbiota during the birthing process [68]. However, recent investigations have posited that the amniotic fluid harbors a resident microbiota, which functions as a primary source of microorganisms for initial colonization of the offspring *in utero* [69–77]. These juxtaposed views have sparked much debate [78–83].

Investigations of human amniotic fluid in normal pregnancy have yielded contradictory results. Multiple studies using DNA sequencing techniques [72, 75–77, 84, 85] and/or quantitative real-time PCR [70, 76] have identified an amniotic fluid microbiota; however, only one of these studies has demonstrated viable microorganisms from amniotic fluid through culture [72] **(Table 1)**. To date, no study has used cultivation, qPCR, and DNA sequencing concurrently to confirm microbial presence in human amniotic fluid during normal pregnancy. The concurrent use of multiple microbiological techniques in such investigations is important because a molecular signal of microorganisms is not necessarily equivalent to a true and viable microbiota [68, 70, 86–88]. For instance, the molecular signal may simply reflect circulating microbial DNA fragments [89]. Furthermore, if there is an amniotic fluid microbiota, it has a very low microbial biomass and, therefore, reliance on molecular techniques such as DNA sequencing to characterize the presumed microbiota is susceptible to influences of background DNA contamination from laboratory environments, DNA extraction kits, PCR reagents, etc. [90]. Yet, very few of the prior investigations that used DNA sequencing techniques to conclude the existence of a human amniotic fluid microbiota incorporated technical controls for background DNA contamination into their analyses [75, 76, 84, 91, 92] **(Table 1)**. Hence, there remains uncertainty as to whether the human amniotic fluid harbors a microbiota.

**Table 1.**
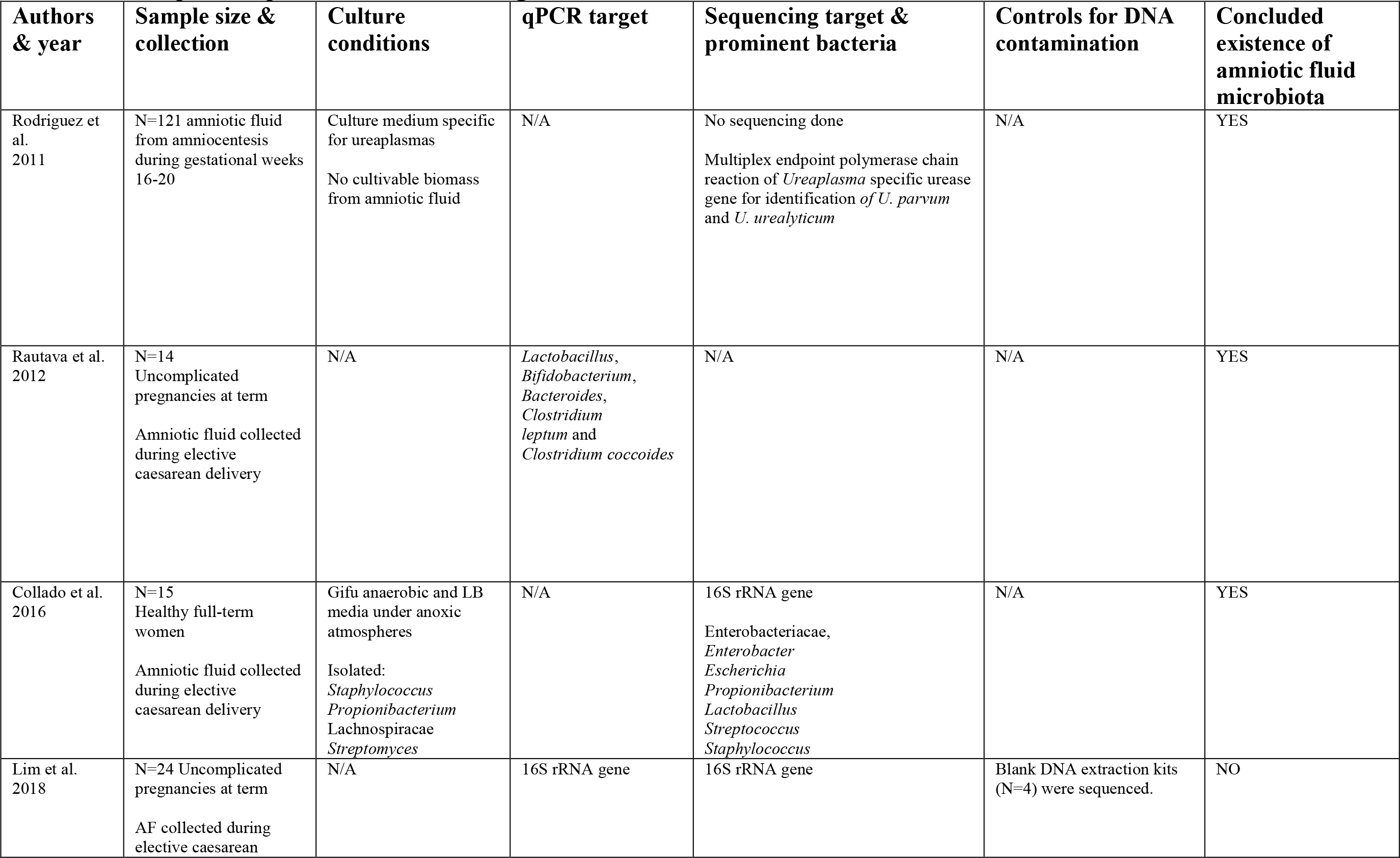
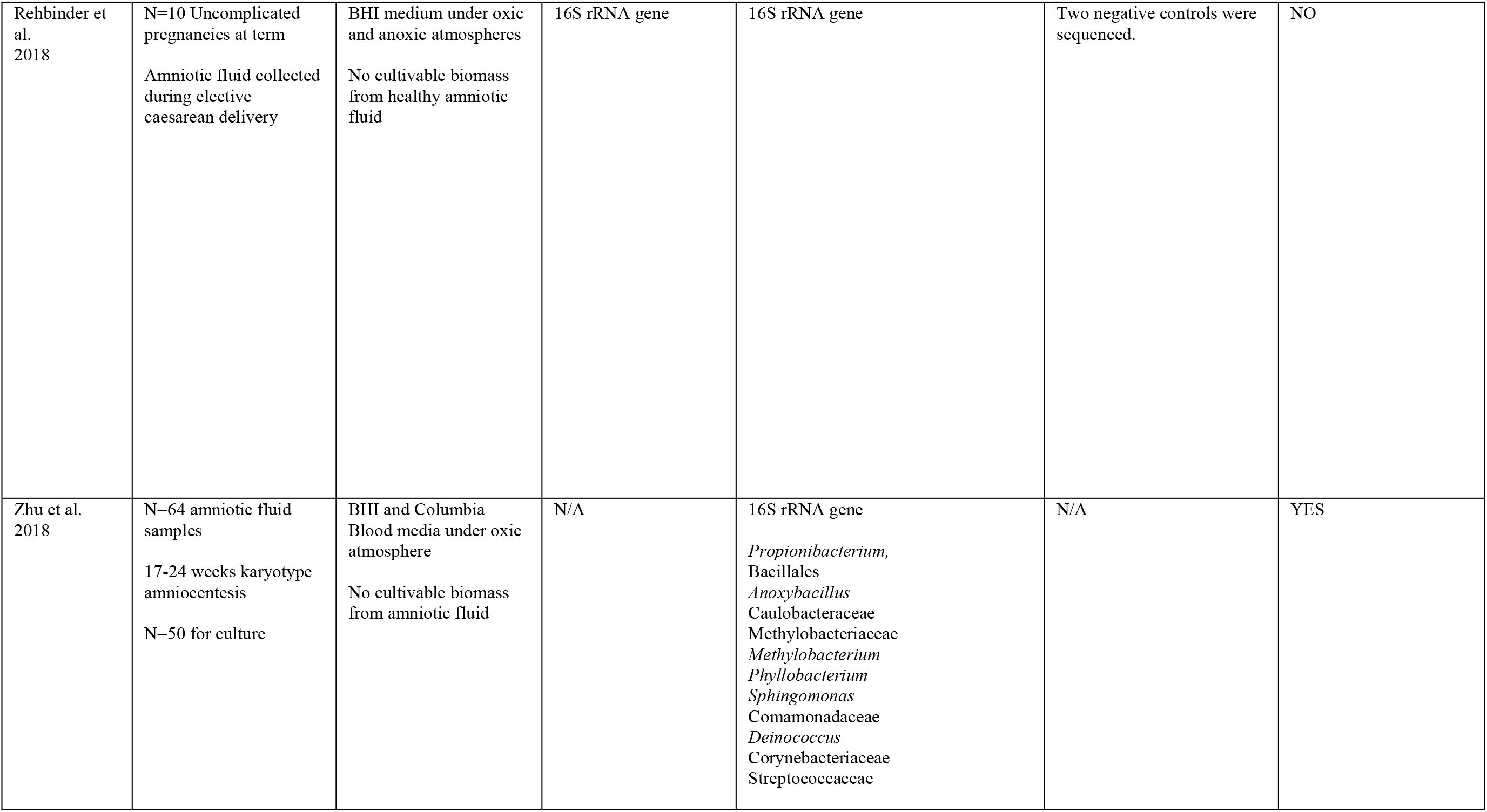
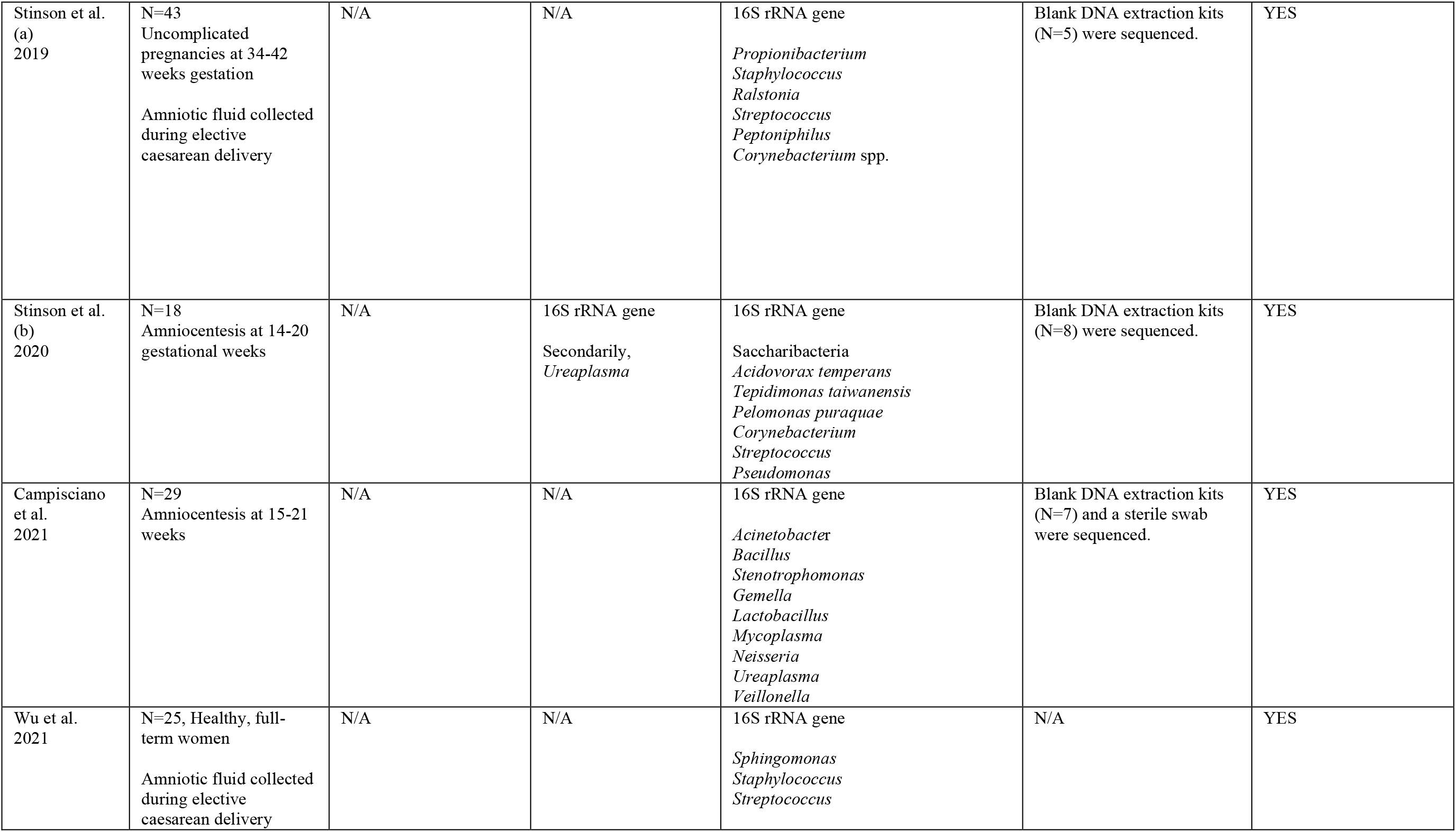
Description of prior molecular investigations of the human amniotic fluid.

The existence of an amniotic fluid microbiota would require a fundamental reconsideration of the role of intra-amniotic microorganisms in fetal development and pregnancy outcomes. Such reconsideration would require the implementation of animal models to perform mechanistic experimentation of host immune-microbe interactions. Yet, there have been only a limited number of studies investigating the presence of an amniotic fluid microbiota in animal models, specifically cattle, horses, sheep, goats, and rats **(Table 2)**. Although each of these studies used DNA sequencing techniques, very few included qPCR, technical controls for background DNA contamination, or culture. Therefore, the objective of the current study was to determine whether the amniotic fluid of mice, the most widely utilized system for studying host immune-microbe interactions [93], harbors a microbiota using technical controls, qPCR, 16S rRNA gene sequencing, and bacterial culture.

**Table 2.**
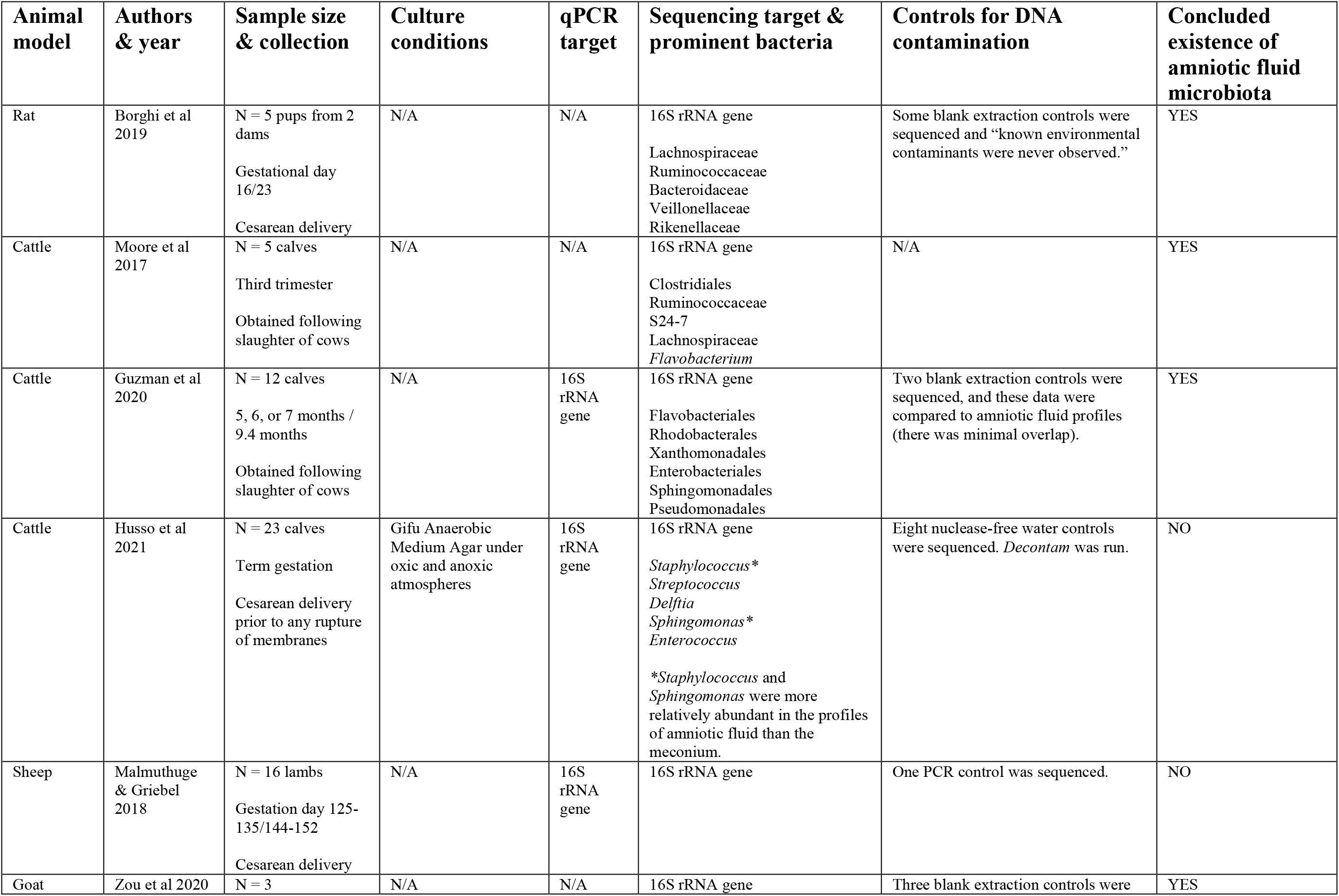
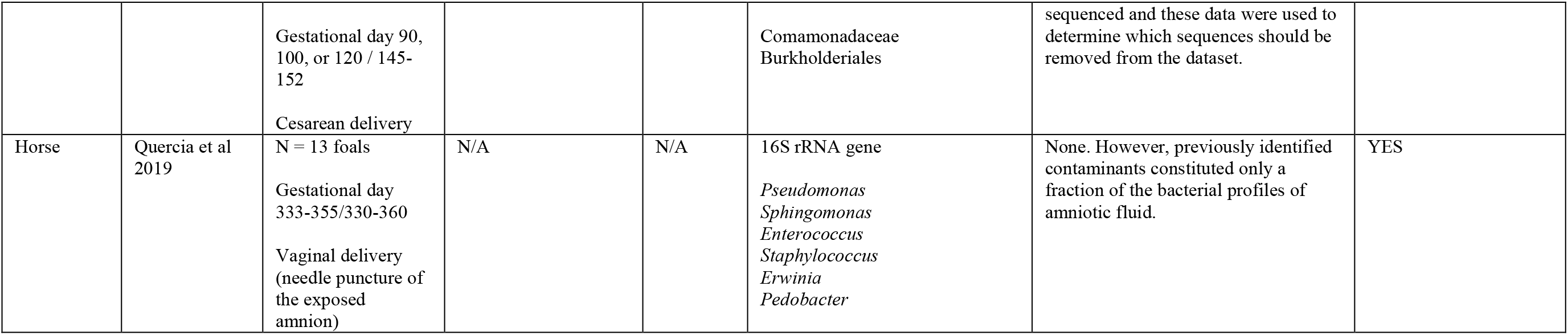
Description of prior molecular investigations of an amniotic fluid microbiota using animal models.

## RESULTS

### Does the murine amniotic fluid contain 16S rDNA?

Amniotic fluid was collected from amniotic sacs located proximally and distally to the cervix under aseptic conditions from 13.5 – 18.5 days *post coitum* (dpc) **(Figure 1)**. First, we evaluated the absolute abundance of 16S rDNA in amniotic fluid using qPCR. There was a significantly higher 16S rDNA signal in proximal (*W* = 6, *p* = 0.0003) and distal (*U* = 16, *p* = 0.004) amniotic fluid samples than in blank extraction controls. However, the 16S rDNA signal did not differ between paired proximal and distal samples (*V* = 89, *p* = 0.571) **(Figure 2A)**. These results indicate that the murine amniotic fluid contains 16S rDNA, and that its concentrations do not depend on proximity to the cervix.

**Figure 1.**
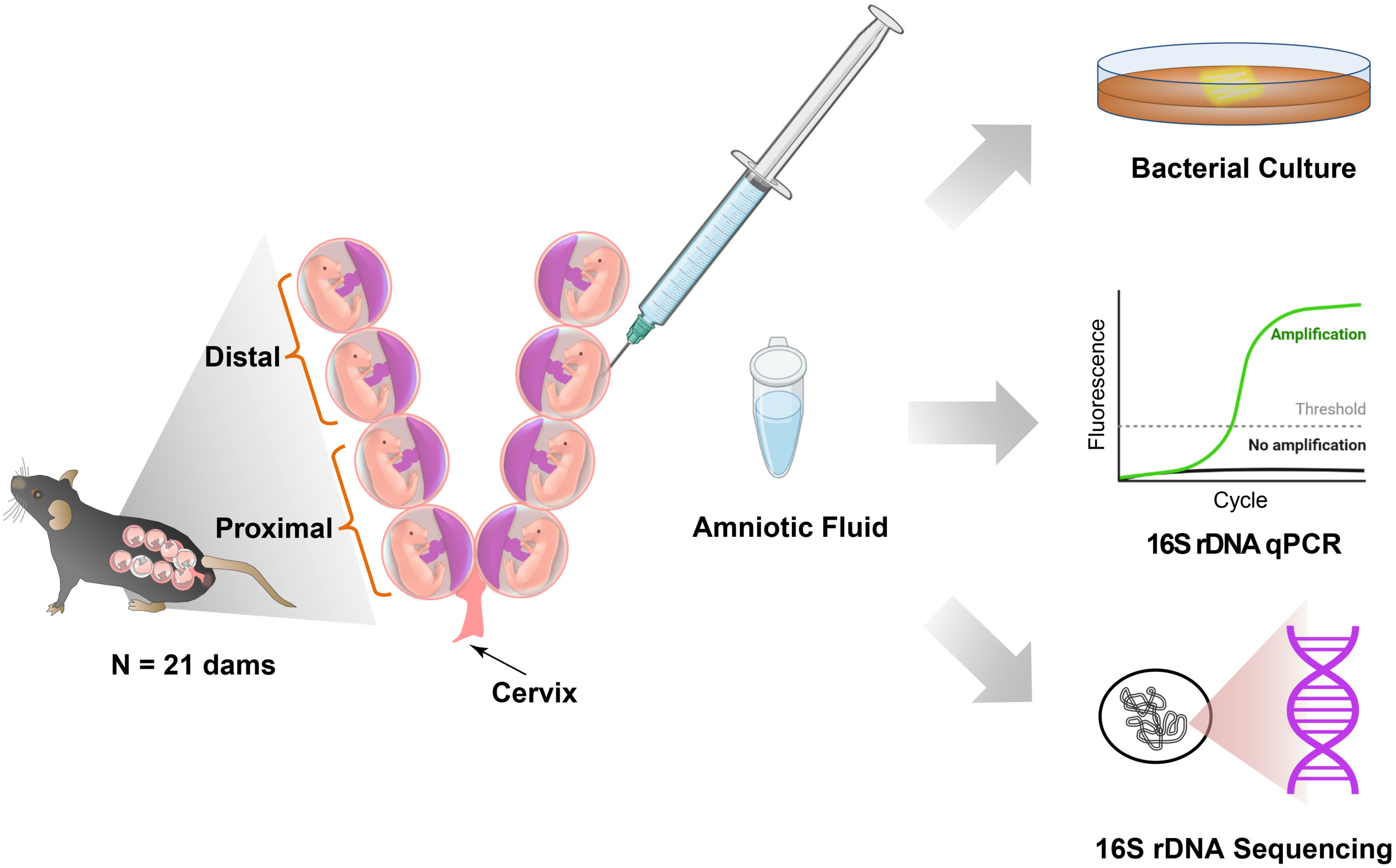
Study design to test for the presence of bacteria in murine amniotic fluid.

**Figure 2.**
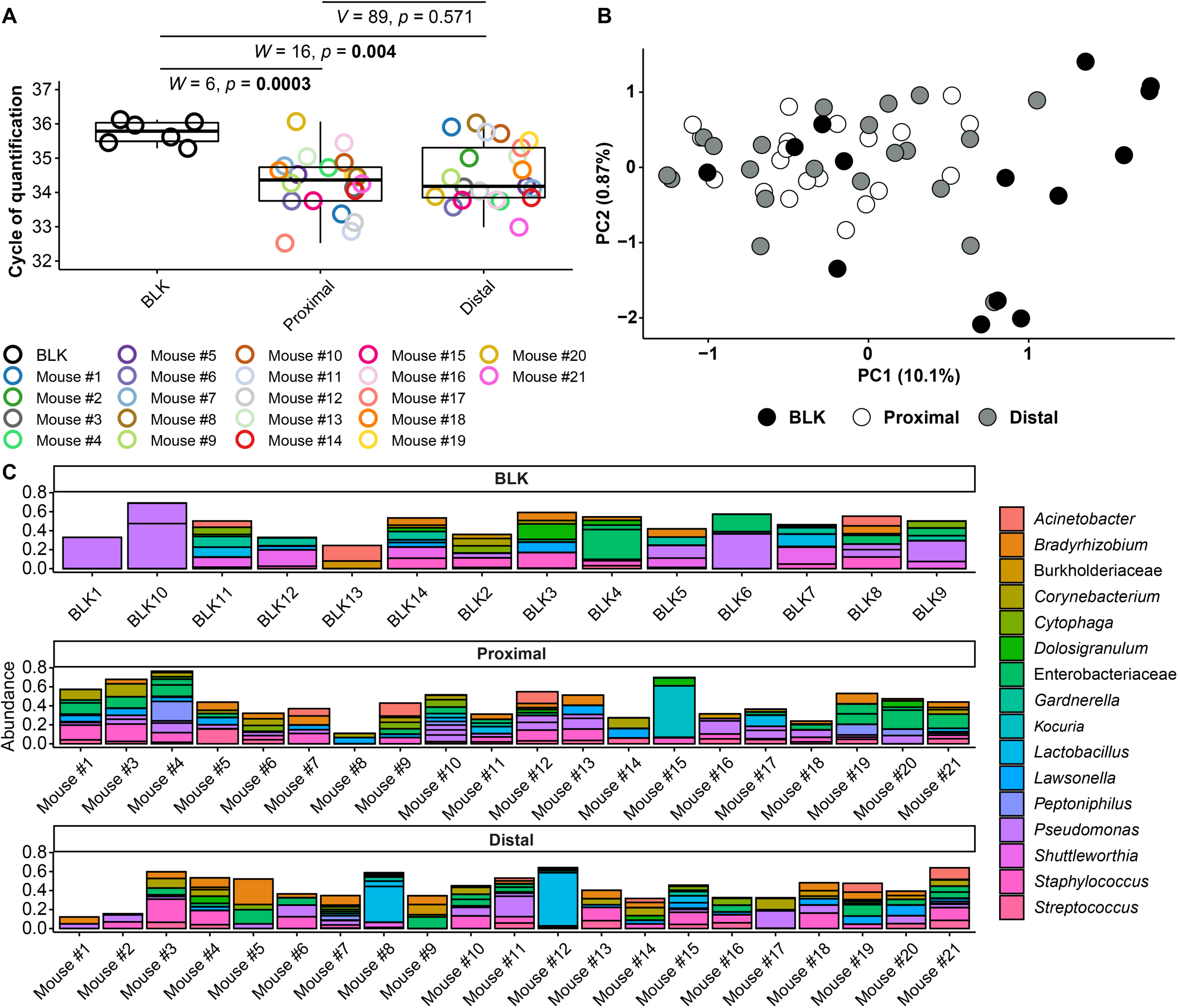
16S rDNA qPCR and sequencing results for amniotic fluid and blank control samples. (A) Cq values from qPCR of proximal and distal amniotic fluid and blank control (BLK) samples. (B) Principal coordinate analysis (PCoA) illustrating variation in 16S rRNA gene profiles among proximal and distal amniotic fluid and blank control samples. The 16S rRNA gene profiles were characterized using the Bray-Curtis similarity index. (C) Taxonomic classifications of the 20 amplicon sequence variants with highest relative abundance across all proximal and distal amniotic fluid and blank control samples.

### Does the 16S rDNA profile differ between murine amniotic fluid and controls?

Next, the 16S rDNA profiles of the amniotic fluid samples were characterized using nucleotide sequencing and the generation of amplicon sequence variants (ASVs). Prior to removing potential contaminants, the 16S rDNA profiles of both the proximal and distal amniotic fluid samples differed from that of negative controls (PERMANOVA *F* = 2.343, *R*^2^ = 0.068, *p* = 0.0001 and *F* = 1.806, *R*^2^ = 0.052, *p* = 0.008, respectively) **(Figure 2B)**. The most prominent ASVs in the proximal and distal amniotic fluid samples and technical controls were *Staphylococcus*, *Pseudomonas*, and *Enterobacteriaceae* (ASVs 4, 6, and 7, respectively) **(Figure 2C)**. Nevertheless, there were differentially abundant taxa between the amniotic fluid samples and negative controls (**Figure 3A and 3B**). Specifically, multiple ASVs classified as *Corynebacterium* were more abundant in proximal (ASV 10) and distal (ASVs 10, 31 and 572) amniotic fluid samples than in controls **(Figure 3A and 3B)**. These corynebacteria were most closely related to *C. tuberculostearicum*, *C*. *mucifaciens*, *C*. *ureicelerivorans*, *C*. *ihumii*, and C. *pilbarense* **(Figure 3C)**. Additional taxa that were differentially abundant in proximal amniotic fluid samples compared to controls were *Streptococcus* (ASV 13) *Pseudomonas* (ASV 24), and *Sphingobium* (ASV 33) **(Figure 3A)**. However, these signals may still represent contamination.

**Figure 3.**
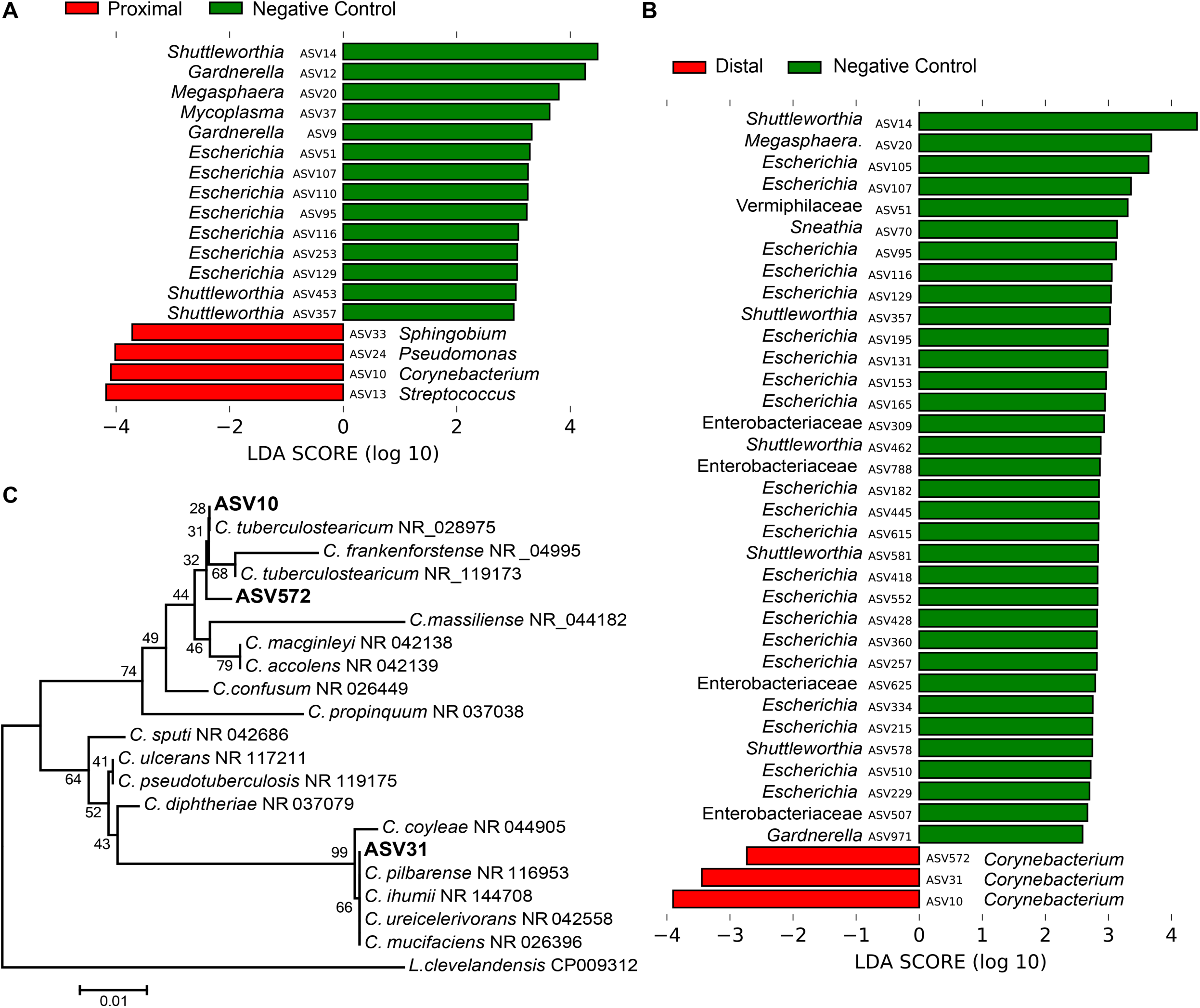
Differentially abundant amplicon sequence variants (ASVs) in proximal and distal amniotic fluid and blank control samples. (A) proximal and (B) distal amniotic fluid samples compared to blank DNA extraction control samples as determined by Linear discriminant analysis effect size analyses. (C) Dendrogram of the three differentially abundant *Corynebacterium* ASVs in amniotic fluid samples and partial 16S rDNA sequences of closely related bacterial type strains. Numbers at the nodes are maximum-likelihood bootstrap values. Scale bar indicates the number of nucleotide substitutions per site.

To address potential background DNA contamination, the program *decontam* was used in part to identify and remove likely contaminants. After contaminants were removed from the dataset, the ASVs with the highest mean relative abundance in both proximal and distal amniotic fluid samples were *Corynebacterium* and *Streptococcus* (ASVs 10 and 13, respectively) **(Figure 4A)**. This is in contrast to the profile structure before contaminant removal **(Figure 2C)**. The 16S rDNA profiles of paired proximal and distal amniotic fluid samples did not differ in richness (Chao1 richness) (*V* = 58, *p* = 0.083) or in evenness (Shannon-Wiener diversity) (*V* = 76, *p* = 0.294). The structure of these profiles did not differ either by mouse ID (PERMANOVA *F* = 0.992, *R*^2^ = 0.495, *p* = 0.551) or proximity to the cervix (*F* = 1.215, *R*^2^ = 0.030, *p* = 0.089) **(Figure 4B)**. Collectively, these results indicate that, if there is a murine amniotic fluid microbiota, it is largely comprised of *Corynebacterium* and *Streptococcus*, both of which are readily grown on brain heart infusion media [94, 95].

**Figure 4.**
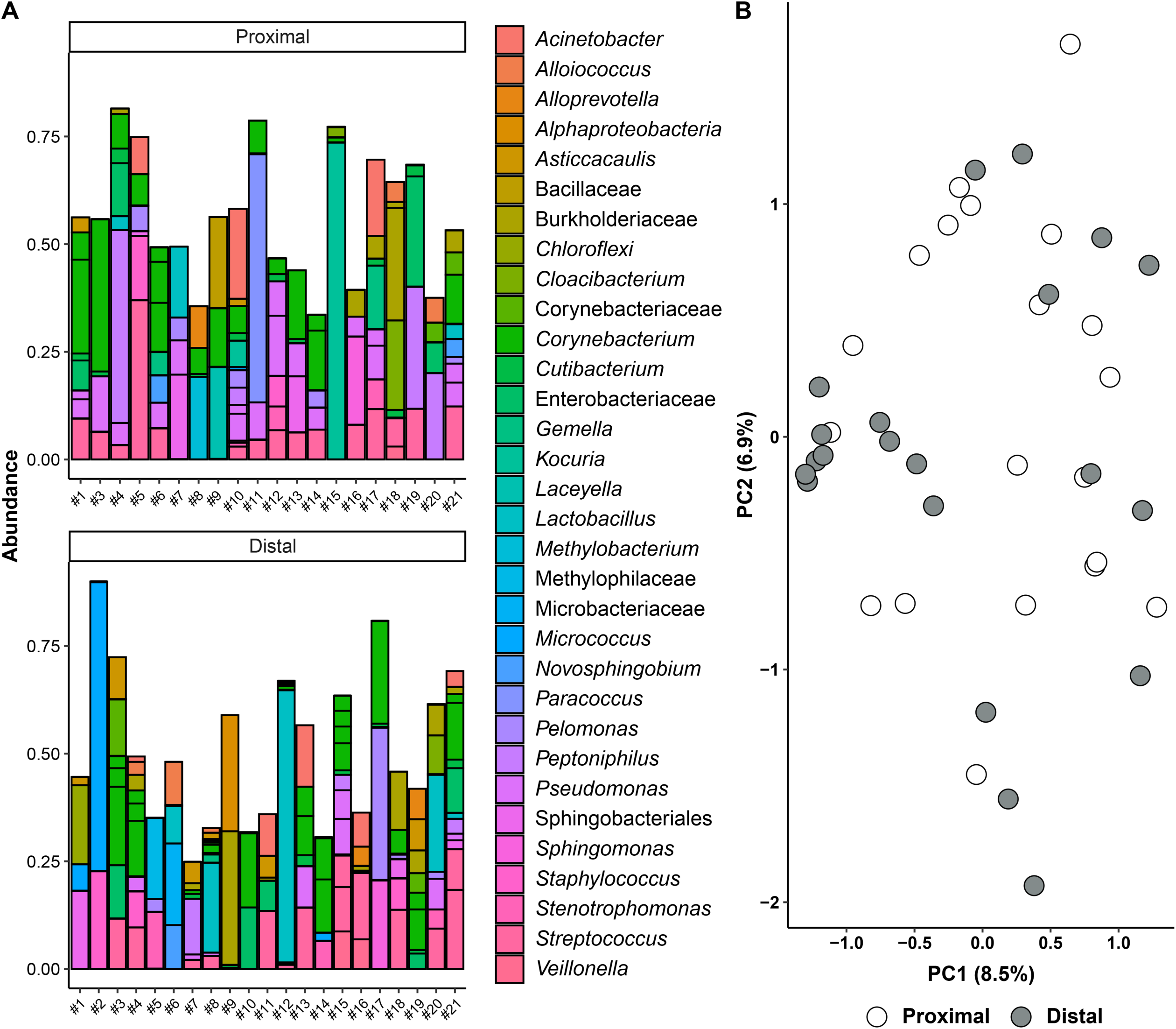
Amniotic fluid sequencing results after the removal of likely contaminating sequences. (A) Bar graph showing the taxonomy of the 45 amplicon sequence variants with highest relative abundance across all proximal and distal amniotic samples. (B) Principal coordinate analysis (PCoA) illustrating variation in 16S rRNA gene profiles among proximal and distal amniotic fluid samples. The 16S rRNA gene profiles were characterized using the Bray-Curtis similarity index.

### Does the murine amniotic fluid contain a viable microbiota?

Forty-two amniotic fluid samples were cultured for bacteria, and only one amniotic fluid sample (Mouse #3 distal) yielded bacterial growth **(Figure 5A)**. For this sample, multiple colonies of a single bacterial morphotype (Gram positive rod) were ultimately recovered under oxic and anoxic conditions. The partial 16S rRNA genes (703 bp) of these isolates were at least 99.7% identical to *Lactobacillus murinus* NBRC 14221 (NR_112689). The proximal and distal amniotic fluid samples from Mouse #3 did not have 16S rDNA concentrations outside the range of other amniotic fluid samples in the study **(Figure 2A)**.

**Figure 5.**
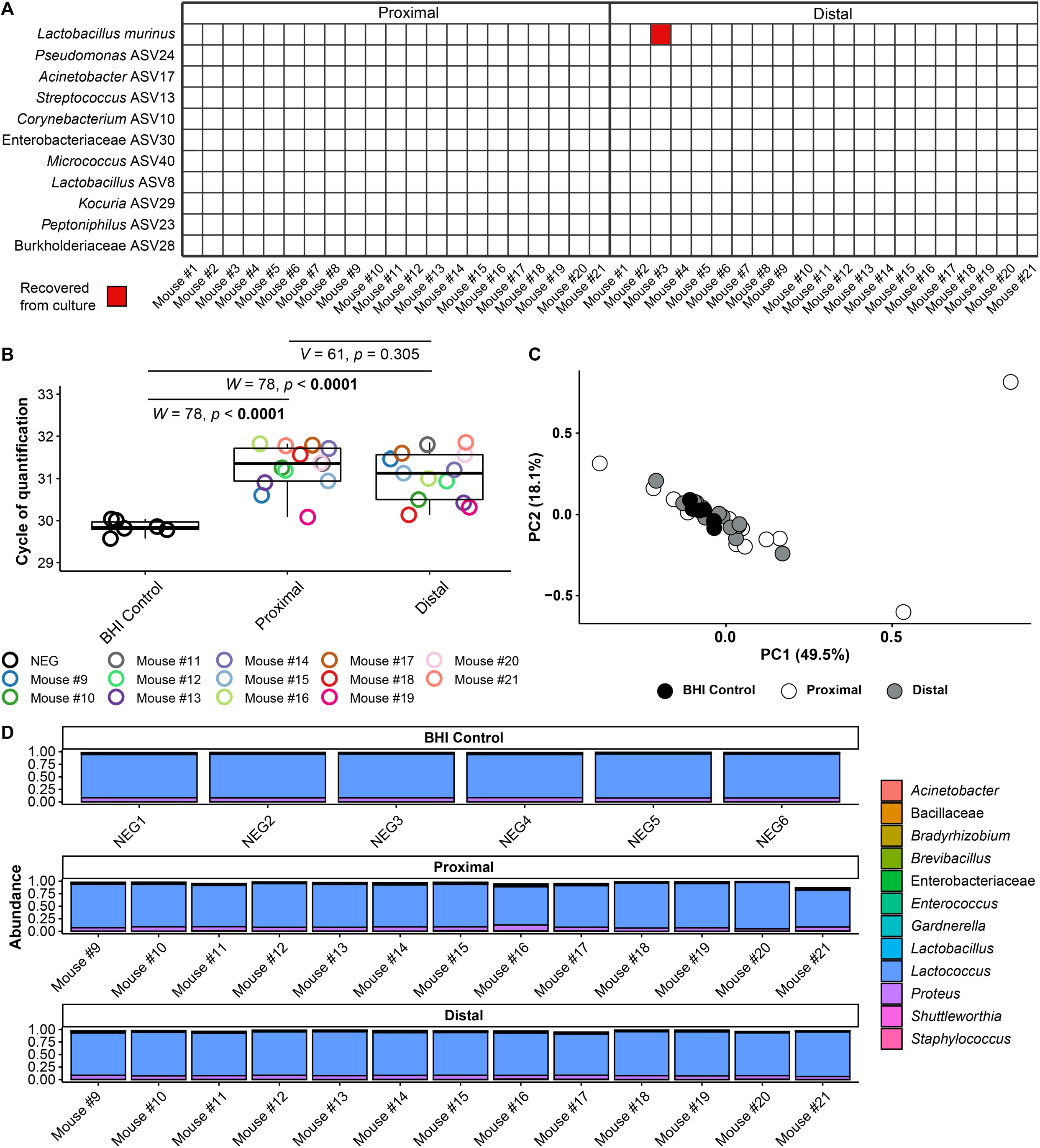
Amniotic fluid culture and blank control 16S rRNA gene qPCR and sequencing results. (A) Bacterial cultivation results for proximal and distal amniotic fluid samples. (B) Cycle of quantification values from qPCR on amniotic fluid culture samples and BHI culture medium controls. (C) Principal coordinate analysis (PCoA) of bacterial relative abundance data from amniotic fluid samples and BHI culture medium controls. (D) Relative abundance of bacteria in the 16S rRNA gene profiles of amniotic fluid samples and BHI culture medium controls.

Secondarily, for 13/21 mice for which culture was attempted, we characterized the 16S rDNA concentration and profile of the amniotic fluid-inoculated BHI broths, and compared these data to those of stock control broth. Overall, the 16S rDNA signal of inoculated broth did not exceed that of stock controls **(Figure 5B)**. Additionally, the 16S rDNA profile structure of both the proximal and distal amniotic fluid cultures did not differ from those of the stock BHI control samples (PERMANOVA *F* = 0.702, *R*^2^ = 0.04, *p* = 0.602 and *F* = 0.918, *R*^2^ = 0.051, *p* = 0.461, respectively) **(Figure 5C and 5D).** Similar to the data for 16S rDNA concentration **(Figure 5B)**, 16S rDNA profile did not differ between paired proximal and distal amniotic fluid samples **(Figure 5C and 5D)**. However, many of these data from sequenced amniotic fluid culture samples may be DNA contaminants.

After removal of contaminants from the dataset using *decontam*, only half of the paired amniotic fluid culture samples (N = 7) had at least 500 sequence reads remaining. The structures of the proximal and distal culture 16S rDNA profiles did not vary by mouse ID (PERMANOVA *F* = 0.815, *R*^2^ = 0.409 *p* = 0.807) or differ based on proximity to the cervix (*F* = 1.057, *R*^2^ = 0.089, *p* = 0.317).

Taken together, using culture and molecular interrogation of culture broths, these data provide no evidence of bacterial growth in proximal or distal amniotic fluids.

## DISCUSSION

In the current study, we utilized quantitative PCR, 16S rRNA gene sequencing, and bacterial culture to investigate the presence of bacterial signals in murine amniotic fluids. Molecular techniques indicated the presence of a 16S rDNA signal in the amniotic fluids; yet, this signal was not verified through culture as coming from a viable microbiota.

### Prior reports of an amniotic fluid microbiota in normal human pregnancy

Investigations using quantitative PCR, 16S rRNA gene sequencing, or cultivation to determine the presence of a human amniotic fluid microbiota in normal pregnancy have yielded inconsistent findings [70, 72, 75–77, 84, 85, 91, 92, 96]. This is likely due in part to insufficient methods such as a lack of multiple complementary techniques for bacterial detection and isolation and/or a lack of appropriate technical controls. Notably, of these studies, only one reported the isolation of bacteria from human amniotic fluid of women who delivered a term neonate [72]. The bacteria that were isolated were *Propionibacterium (Cutibacterium)* and *Staphylococcus.* These bacteria were also identified in the 16S rRNA gene profiles of amniotic fluid; however, these bacteria are typical inhabitants of the human skin and may therefore represent skin contaminants [97].

Overall, of the studies that have performed 16S rRNA gene sequencing to investigate the existence of a human amniotic fluid microbiota in normal pregnancies [72, 75–77, 84, 85, 91, 92, 96], only five included technical controls for background DNA contamination [75, 76, 84, 91, 92]. Three concluded the existence of an amniotic fluid microbiota, although these studies did not include a culture component [75, 76, 84]. The first study [75] reported that 83.7% (36/43) of amniotic fluid samples had a 16S rDNA signal, with varying degrees of *Propionibacterium* (*Cutibacterium*) *acnes*, *Staphylococcus epidermidis, Ralstonia*, *Streptococcus anginosus*, and *Peptoniphilus* dominance. The second study [76] reported that 19.9% (238/1,206) of amniotic fluid samples yielded a 16S rDNA signal; they were dominated by Saccharibacteria, *Acidovorax*, *Tepidimonas*, *Pelomonas,* and *Streptococcus oligofermentans*. In the third study [84], only 13.8% (4/29) of amniotic fluid samples had a detectable 16S rDNA signal, with *Actinomyces*, *Cutibacterium*, *Staphylococcus*, and *Streptococcus* being most relatively abundant. Thus, the most reported bacterial taxa detected in human amniotic fluid investigations were *Staphylococcus* and *Cutibacterium*, two typical skin bacteria [97]. These results illustrate the need for more comprehensive investigations using multiple complementary modes of microbiologic inquiry, as well as the need for appropriate technical controls.

### Existence of an amniotic fluid microbiota in animal models

In cattle, three investigations utilized 16S rRNA gene sequencing to explore the presence of an amniotic fluid microbiota [98–100] (**Table 2**). Two concluded the existence of an amniotic fluid microbiota using this approach [98, 99]; however, one study, which also included qPCR and culture, concluded that the bacterial signals in the amniotic fluid did not exceed those in controls [100]. In two investigations of horses and goats, a microbiota was identified in the amniotic fluid using 16S rRNA gene sequencing [101, 102]. However, in a study of sheep, the amniotic fluid was determined to be sterile using qPCR and 16S rRNA gene sequencing [103].

In the only study to date of rodents [73], 16S rRNA gene sequencing was used to demonstrate that amniotic fluid microbiota profiles were pup- and dam-specific in a rat model, yet they were not different from those of the placenta or fetal intestine. The primary bacteria detected were identified as Lachnospiraceae, Ruminococcaceae, Bacteroidaceae, Veillonellaceae, Rikenellaceae, and Proprionibacteriaceae [73]. However, this study did not include qPCR or culture components.

### Our findings in the context of prior studies

In the current study, quantitative PCR showed significantly greater 16S rRNA gene signal in both proximal and distal amniotic fluid samples than in the negative controls, indicating the presence of 16S rDNA in amniotic fluid samples regardless of proximity to the cervix. These findings are consistent with the qPCR results of a prior study of cattle amniotic fluid [99].

Our investigation using 16S rRNA gene sequencing detected higher relative abundances of DNA from *Corynebacterium* spp., *Pseudomonas*, *Sphingobium*, and *Streptococcus* in the amniotic fluid of mice than in controls (**Figure 3)**. *Corynebacterium* spp. and *Streptococcus* spp. are resident microbiota of mammals, including humans and mice [97, 104–106]. However, these microorganisms have also been identified as common bacterial DNA contaminants in studies with low microbial biomass [84, 90]. *Corynebacterium* spp. are aerobic, non-spore-forming, Gram-positive bacteria [94] that have been identified as members of the mouse skin [106] and respiratory [105] microbiotas. Specifically, *Corynebacterium tuberculostearicum* (ASV 10) has been previously detected in human amniotic fluid using molecular techniques; however, this bacterium was not recovered using conventional culture methods [75, 107]. The *Streptococcus* ASV detected in the current study (ASV 13) had an identical sequence match with multiple members of the Mitis group of the genus *Streptococcus,* which are common inhabitants of the oral cavity and upper respiratory tract in humans [108] and have been detected in the lungs of mice [109]. *Pseudomonas* is widely distributed amongst mammals and the broader environment [110]. In our study, BLAST analysis was performed on ASV 24 (*Pseudomonas*), but a species-level taxonomy could not be assigned, indicating that the V4 region of the 16S rRNA gene is not adequate for differentiation of *Pseudomonas* species. *Sphingobium* is typically an environmental microorganism [111]. In the current study, BLAST analysis for ASV 33 showed that it was identical to the typical soil bacteria *S. naphthae*, *S. olei*, and *S. soli* [112–114]. A single case was reported of *S. olei* causing peritonitis via infection of an indwelling peritoneal catheter in a patient with end stage renal disease [115]. In summary, although some of these microorganisms have been found in biologically relevant sites, the importance of their DNA signal in amniotic fluid in this study requires further investigation.

An inherent limitation of molecular investigations is the inability to differentiate between whether the presence of 16S rDNA signal is due to the presence of viable bacteria, dead cells, or environmental DNA. While many studies have used molecular techniques to confirm the existence of bacterial DNA in the placenta, fetal tissue, and amniotic fluid [48, 70, 72, 75–77, 84, 85, 91, 92, 96], only some have attempted to culture bacteria from these same samples [72, 85, 92, 96]. Notably, *Corynebacterium*, *Pseudomonas*, *Sphingobium, Streptococcus,* and other prominent bacteria identified in molecular surveys were not recovered in culture in this study. Indeed, the only microorganism that was cultured, *Lactobacillus murinus*, was not detected in the 16S rRNA gene profile of any amniotic fluid sample. *L*. *murinus* is known to reside in the GI system of mice, where it has been documented to play a role in attenuating inflammation [116]. Indeed, in a prior study [109], *L. murinus* was found in multiple body sites of pregnant mice. Given its wide distribution among and within mice, this *Lactobacillus* isolate may represent a culture contaminant.

### Strengths of this study

The current study has three principal strengths. First, we used multiple, complementary modes of inquiry, including 16S rRNA gene qPCR, 16S rRNA gene sequencing, and bacterial culture to assess whether there is an amniotic fluid microbiota in mice. Furthermore, the culture component of the study included molecular validation. Second, we utilized robust sterile techniques as well as negative, experimental, and positive controls when performing extractions and molecular work to assure that any bacterial DNA signal detected in the experimental samples could be correctly attributed to a true 16S rDNA signal in the amniotic fluid versus environmental or reagent contamination. Third, we sampled amniotic fluid from amniotic sacs proximal and distal to the cervix for assessing differential presence of microorganisms throughout the uterine horns of mice.

### Limitations of this study

The current study has two principal limitations. First, this study focused exclusively on assessing the presence of bacteria in murine amniotic fluid, whereas viruses and eukaryotic microorganisms were not considered in this study. Second, we used a specific animal model and therefore interpretation of results should consider the potential effect of variation in physiological and morphological characteristics among mouse strains and across animal facilities.

### Conclusion

Using qPCR, 16S rRNA gene sequencing, and bacterial culture, we did not find consistent or reproducible evidence of an amniotic fluid microbiota in mice. This study provides evidence against amniotic fluid as a source of microorganisms for colonization of the fetus, and illustrates the importance of using multiple methodologies and the appropriate technical controls in investigations assessing microbial profiles of body sites historically presumed to be sterile.

## MATERIALS AND METHODS

### Study subjects

C57BL/6 mice were obtained from The Jackson Laboratory (Bar Harbor, ME, USA) and bred at the C.S. Mott Center for Human Growth and Development at Wayne State University, Detroit, MI, USA in the specific-pathogen-free (SPF) animal care facility. Mice were housed under a 12 h:12 h light/dark cycle and had access to food (PicoLab laboratory rodent diet 5L0D; LabDiet, St. Louis, MO, USA) and water *ad libitum*. Females (8-12 weeks old) were mated with males of demonstrated fertility. Daily examination was performed to assess the appearance of a vaginal plug, which indicated 0.5 days *post coitum* (dpc). Dams were then housed separately from the males and their weights were checked daily. An increase in weight of ≥ 2 g by 12.5 dpc confirmed pregnancy. All procedures were approved by the Institutional Animal Care and Use Committee (IACUC) (Protocol No. 18-03-0584).

### Sample collection and storage

Twenty-one pregnant mice were included in this study **(Figure 1)**. Pregnant mice were euthanized during the second half of pregnancy (13.5-18.5 dpc). The abdomen was shaved, and 70% ethanol was applied. Dams were placed on a sterile surgical platform within a biological safety cabinet. Study personnel wore sterile sleeves, masks, and powder-free sterile gloves during sample collection, and sterile disposable scissors and forceps were utilized. Iodine was applied to the abdomen with a sterile cotton swab, and after the iodine dried a midline skin incision was performed along the full length of the abdomen. The peritoneum was longitudinally opened using a new set of scissors and forceps, and the uterine horns were separated from the cervix and placed within a sterile petri dish. A sterile syringe with a 26G needle was utilized to obtain amniotic fluid from amniotic sacs proximal to the cervix and from amniotic sacs that were distal from the cervix. Due to the small volume of amniotic fluid often obtained from each amniotic sac (< 40 μl), amniotic fluid was obtained from two adjoining amniotic sacs and pooled. The amniotic fluid was aliquotted into two sterile tubes and transported immediately to the microbiology lab for bacterial culture and molecular analyses, respectively. The tube with the amniotic fluid for molecular analyses was stored at −80°C.

### Culture of amniotic fluid samples

For all mice, proximal and distal amniotic fluid samples (~40 μL each) were cultured in 200 μL Brain-Heart-Infusion (BHI) broth supplemented with 5 mg/L of hemin and 2 μg/L of vitamin K under oxic and anoxic conditions for 48 hours. For the first eight mice in the study, 40 μL of the BHI culture was then plated on supplemented BHI agar plates and cultured under the respective atmospheric condition for an additional 48 hours, and resultant bacterial isolates were taxonomically characterized. For the last 13 mice in the study, 40 μL of the BHI culture was subsequently plated on supplemented BHI agar plates and cultured under the respective atmospheric condition if turbidity of the broth culture was observed after 48 hours of incubation. Any potential growth of bacteria in BHI broth cultures of proximal and distal amniotic fluid samples was then assessed through qPCR and 16S rRNA gene sequencing. As each amniotic fluid sample was cultured under oxic and anoxic conditions, 125 μL each from the oxic and anoxic broth cultures were pooled and stored at −80°C. The 16S rRNA gene loads and profiles of these amniotic fluid broth cultures were compared to those of six uninoculated BHI broth negative controls using qPCR and 16S rRNA gene sequencing.

### DNA extraction

Genomic DNA was extracted within a biological safety cabinet from amniotic fluid and BHI broth samples, as well as positive (i.e., human clean catch urine (N=3) and negative (i.e., human amniotic fluid (N=3), sterile BHI broth (N=6), blank DNA extraction kits (N=14)) controls using the DNeasy PowerLyzer Powersoil kit (Qiagen, Germamtown, MD, USA), with minor modifications to the manufacturer’s protocols as previously described [109, 117]. Specifically, following UV treatment, 400 μL of Powerbead solution, 200 μL of phenol:chloroform:isoamyl alcohol (pH 7-8), and 60 μL of preheated solution C1 were added to the provided bead tubes. Next, 250 μL amniotic fluid or BHI sample were added to the tubes. When less than 250 μL of amniotic fluid was available (9/41 samples, 21%) a minimum of 100 μL was added. Tubes were briefly vortexed and cells were then mechanically lysed in a bead beater for two rounds of 30 sec each. Following 1 minute of centrifugation, supernatant was transferred to new tubes and 1 μL of PureLink™ RNase A (20mg/mL, Invitrogen), 100 μL of solution C2, and 100 μL of solution C3 were added. Tubes were then incubated at 4°C for 5 min. After a 1 min centrifugation, lysates were transferred to new tubes containing 650 μL of C4 solution and 650 μL of 100% ethanol. Lysates were then loaded onto filter columns 635 μL at a time, centrifuged for 1 min, and the flowthrough discarded. This wash process was repeated three times to ensure all lysate passed through the filter columns. Following the wash steps, 500 μL of solution C5 was added to the filter columns and centrifuged for 1 min. After discarding the flowthrough, the tubes were centrifuged for 2 min to dry the filter columns. The spin columns were transferred to clean 2.0 mL collection tubes and 60 μL of pre-heated solution C6 was added directly to the center of the spin columns. Following a 5 min room temperature incubation, DNA was eluted by centrifuging for 1 min. Purified DNA was then transferred to new 2.0 mL collection tubes and stored at −20°C.

### 16S rRNA gene quantitative real-time PCR

To measure total 16S rRNA gene abundance within samples, amplification of the V1-V2 region of the 16S rRNA gene was performed according to the protocol of Dickson et al. [118], with minor modifications as previously described [109, 117]. The modifications consisted of using a degenerative forward primer (27f-CM: 5’-AGA GTT TGA TCM TGG CTC AG-3’) and degenerate probe with locked nucleic acids (+) (BSR65/17: [5’-56FAM-TAA +YA+C ATG +CA+A GT+C GA-BHQ1-3’]). Each 20 μL reaction was performed with 0.6 μM of 27f-CM primer, 0.6 μM of 357R primer (5’-CTG CTG CCT YCC GTA G-3’), 0.25 μM of BSR65/17 probe, 10.0 μL of 2X TaqMan Environmental Master Mix 2.0 (Invitrogen), and 3.0 μL of purified DNA or nuclease-free water. The following conditions were used to perform the total bacterial DNA qPCR: 95° C for 10 min, and then 40 cycles of 94°C for 30 sec, 50°C for 30 sec, and 72°C for 30 sec. Each reaction was performed in triplicate using an ABI &500 thermocycler (Applied Biosystems, Foster City, CA, USA). After normalization to the ROX passive reference dye, the 7500 Software version 2.3 (Applied Biosystems, Foster City, CA, USA) was used to analyze the raw amplification data with the default threshold and baseline settings. Calculation of the cycle of quantification (Cq) values for the samples was based upon the mean number of cycles necessary for the exponential increase of normalized fluorescence.

### 16S rRNA gene sequencing

The V4 region of the 16S rRNA gene was amplified and sequenced via the dual indexing strategy developed by Kozich et al. [119]. The forward and reverse primers used were 515F: 5’-GTGCCAGCMGCCGCGGTAA-3’ and 806R: 5’-GGACTACHVGGGTWTCTAAT-3’, respectively. Duplicate 20 μL PCR reactions were performed containing 0.75 μM of each primer, 3.0 μL DNA template, 10.0 μL of DreamTaq High Sensitivity Master Mix (Thermo Scientific, Waltham, MA, USA), and 5 μL of DNase-free water. Reaction conditions were as follows: 95° for 3 min, followed by 38 cycles of 95°C for 45 sec, 50°C for 60 sec, and 72°C for 90 sec, followed by an additional elongation at 72°C for 10 min. The duplicate PCR reactions were then pooled, and DNA was quantified with a Qubit 3.0 fluorometer and Qubit dsDNA assay kit (Life Technologies, Carlsbad, CA) following the manufacturer’s protocol. Samples were pooled in equimolar concentrations and purified using the Cytiva Sera-Mag Select DNA Size Selection and PCR Clean-Up Kit (Global Life Sciences, Little Chalfont, Buckinghamshire, UK) according to the manufacturer’s instructions.

The R package *decontam* version 1.6.0 [120] was used to identify ASVs that were likely potential background DNA contaminants based on their distribution among biological samples (amniotic fluid and BHI cultures) and negative controls (blank DNA extractions and stock BHI broth) using the “IsNotContaminant” method. Identification of contaminant ASVs was assessed for amniotic fluid and BHI cultures independently. An ASV was determined to be a contaminant, and was removed from the dataset, if it had a *decontam* P score ≥ 0.7 and was present in at least 20% of negative controls with an overall average relative abundance of at least 1.0%

### Statistical analysis

Prior to statistical analyses, the bacterial profiles of proximal and distal amniotic fluid samples and blank DNA extraction controls were rarefied to 1,366 sequence reads (set.seed = 1) using phyloseq [121]. The bacterial profiles of proximal and distal BHI culture samples and stock BHI broth samples were rarefied to 21,227 sequence reads. Variation in the bacterial profiles was visualized through Principal Coordinates Analyses (PCoA) using the R package vegan version 2.5-6 [122]. Alpha diversity values and 16S rDNA signal (qPCR Cq) values across sample groups were compared using the “wilcox.test” function in R version 3.6.0 [123]. Beta diversity of amniotic fluid bacterial profiles was characterized using the Bray-Curtis dissimilarity index. Bacterial community structure of amniotic fluid and BHI culture samples was compared using PERMANOVA [124] with the “adonis” function in the R package vegan version 2.5-6 [122]. Assessment of differentially abundant taxa across sample groups was performed using Linear Discriminant Analysis Effect Size, or LEfSe [125] with default parameters. Analysis of the phylogenetic relationships of selected ASVs and other bacteria was performed using the Neighbor-Joining method [126] in MEGA 6 software [127] with the Maximum Composite Likelihood method and bootstrapping of 1,000 replicates, allowing for transitions and transversions.

## DATA AVAILABILITY

Sample-specific MiSeq run files have been deposited on the NCBI Sequence Read Archive (BioProject ID PRJNA751620).

